# General remarks of the secondary promoter element regulating the genome replication of paramyxo- and filoviruses

**DOI:** 10.1101/2023.11.26.568702

**Authors:** Shoichi Ashida, Shohei Kojima, Takashi Okura, Fumihiro Kato, Shuzo Urata, Yusuke Matsumoto

**Affiliations:** Transboundary Animal Diseases Research Center, Joint Faculty of Veterinary Medicine, Kagoshima University, Kagoshima, Japan; Genome Immunobiology RIKEN Hakubi Research Team, RIKEN Center for Integrative Medical Sciences, Yokohama, Japan; Department of Virology 3, National Institute of Infectious Diseases, Tokyo, Japan; National Research Center for the Control and Prevention of Infectious Diseases (CCPID), Nagasaki University, Nagasaki, Japan

## Abstract

Paramyxo- and filovirus genomes are equipped with bipartite promoters at their 3’ ends to initiate RNA synthesis. The two elements, the primary promoter element 1 (PE1) and the secondary promoter element 2 (PE2), are separated by a spacer region that must be precisely a multiple of six nucleotides, indicating these viruses are related to the “rule of six”. However, our knowledge of PE2 has been limited to a narrow spectrum of virus species. In this study, a comparative analysis of 1,647 paramyxoviral genomes from a public database revealed that the paramyxovirus PE2 can be clearly categorized into two distinct subcategories: one marked by C repeats at every six bases (exclusive to the subfamily *Orthoparamyxovirinae*), and another characterized by CG repeats every six bases (observed in the subfamilies *Avulavirinae* and *Rubulavirinae*). This unique pattern collectively mirrors the evolutionary lineage of these subfamilies. Furthermore, we showed that the PE2 of the *Rubulavirinae*, with the exception of mumps virus, serves as part of the gene-coding region. This may be due to the fact that the *Rubulavirinae* is the only paramyxovirus that cannot propagate without RNA editing occurring. Zaire ebolavirus has eight sequential uracil (U) repeats every six bases within its genomic promoter. We showed that a minimum of four sequential U-containing hexamer repeats is imperative for genome replication. This discovery led to the identification of such quadruplet U-containing hexamer repeats within the genomic and antigenomic promoters of other viruses within the family *Filoviridae*.

**Significance:** The genomic intricacies of paramyxo- and filoviruses are highlighted by the bipartite promoters—PE1 and PE2—at their 3’ termini. The spacer region between these elements follows the “rule of six”, crucial for genome replication. By a comprehensive analysis of paramyxoviral genome sequences, we identified distinct subcategories of PE2 based on C and CG repeats that were specific to *Orthoparamyxovirinae* and *Avulavirinae*/*Rubulavirinae*, respectively, mirroring their evolutionary lineages. Notably, the PE2 of *Rubulavirinae* is integrated in the gene-coding region, a unique trait potentially linked to its complete dependence on RNA editing for virus growth. This study also focused on the PE2 sequences in filovirus genomes. In filoviruses, four consecutive U-containing hexamer repeats appeared to be critical for their genome replication.

## Introduction

The order *Mononegavirales* is a group of non-segmented negative-strand RNA viruses that includes the families *Rhabdoviridae*, *Paramyxoviridae*, *Pneumoviridae*, *Filoviridae* and *Bornaviridae*, all of which contains many important animal and human pathogens that cause diseases, such as rabies, measles, respiratory syncytial virus disease and Ebola virus disease. All viruses in the order *Mononegavirales* share a common mechanism of mRNA transcription and genome replication (1). The negative-strand RNA genome is entirely encapsidated by viral nucleoprotein (NP), which serves as template for viral RNA-dependent RNA polymerase (RdRp) complex composed of large protein (L) and co-factor (s) such as phosphoprotein (P) or VP35. Although it is common that the viral genomic 3’ terminus acts as a replication promoter (termed promoter element 1; PE1) for viral RdRp, viruses in the families *Paramyxoviridae* and *Filoviridae* possess bipartite promoters, which require a secondary promoter element (PE2) located in the internal genomic region (2–6). The paramyxo- and filoviruses employ an RNA editing mechanism in the P gene and glycoprotein gene, respectively. During the mRNA transcription process, viral RdRp recognizes the cis-acting element of RNA editing signal within each gene. Co-transcriptionally, non-template nucleotides (nts) are appended to the mRNA, enabling the synthesis of multiple proteins from a single gene (7–9). It is important to note that all viruses possessing bipartite promoters and RNA editing are inherently subject to the “rule of six,” which requires hexamer-phasing of nts within a genomic region to facilitate viral propagation (2–6,10). In both paramyxo- and filoviruses, the genomic RNA is covered by NPs at exactly every 6 nts (11–14).

The bipartite replication promoters play an essential role in that the virus genome is recognized as the multiple of six by RdRp of paramyxoviruses. The paramyxoviral PE2 is characterized by the presence of specific nts at every 6 nts in the internal genomic region. Paramyxovirus nucleocapsids have 13 NP subunits per turn such that PE1 and PE2 are juxtaposed on the same face of the nucleocapsid helix for concerted recognition by the viral RdRp (15,16). In Sendai virus (SeV) in the subfamily *Orthoparamyxovirinae*, the 14th to 16th hexamers contain 5’-GAAGA<u>C</u> UUGGA<u>C</u> UUGUC<u>C</u>-3’, in which a C is found at every 6 nts three times (Fig. 1A and B, PE2) (2). In parainfluenza virus type 5 (PIV5) in the subfamily *Rubulavirinae*, the 13 to 15th hexamers contain 5’-<u>CG</u>GGA <u>CG</u>AUGG <u>CG</u>AGGA-3’, in which CG are found at every 6 nts three times (Fig. 1A and B, PE2) (4). If a 1 nt insertion occurs upstream in the genome, the repeated C in SeV and the repeated CG in PIV5 would be shifted by 1 nt (Fig. 1B, 1 nt insertion). In this case, viral RdRp cannot recognize the genome as a correct template, resulting it to non-template for further replication, which acts to keep the remaining genome in a multiple of six in the infected cells. Although the PE2 is thus essential for viral genome replication, its characteristics have only been studied in limited numbers of virus species, and it remains unclear whether these regulatory mechanisms are universal to all known paramyxoviruses.

**Figure 1.**
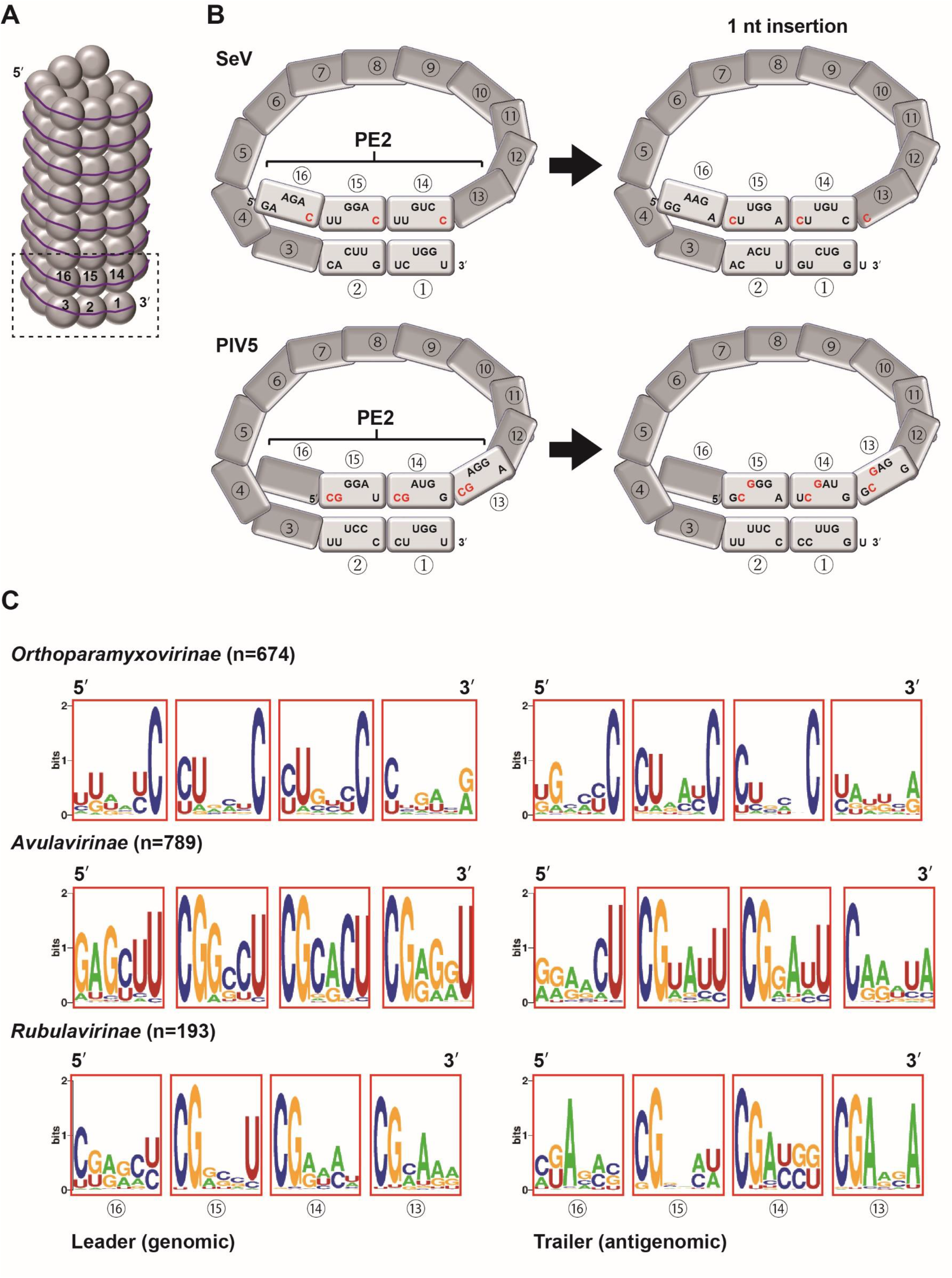
Extensive comparative analysis of the PE2 sequences of viruses in the family *Paramyxoviridae.* (A) A schematic diagram of the paramyxovirus helical nucleocapsid. The dotted box indicates the replication promoters that are shown in more detail in Figure 1B. The gray circles indicate NPs, and the numbers in the circles indicate the number of the NP hexamer from the 3’ terminus. The line indicates RNA. (B) Schematic diagrams of the replication promoters for the genomic RNA of SeV and PIV5. The NPs (gray squares) bind to a sequence of 6 nts. The numbers in the circles indicate the number of the NP hexamer from the 3’ terminus. The normal states are shown on the left, while the nt positions when a single nt insertion has occurred upstream are shown on the right. The 24 nts spanning from the 13th to the 16th NPs from the 3’ terminus collectively constitute the PE2. The conserved nts in PE2 are highlighted in red. (C) Conserved nts within the leader and trailer promoters of viruses belonging to the subfamilies *Orthoparamyxovirinae*, *Avulavirinae* and *Rubulavirinae*. The numbers in the circles on the diagram indicate the number of the hexamer from the 3’ terminus.

The family *Filoviridae* contains several genera, including *Ebolavirus*, *Marburgvirus* and *Cuevavirus*. The genus *Ebolavirus* contains at least six viruses, including Zaire ebolavirus (ZEBOV), Sudan ebolavirus (SUDV), Tai forest ebolavirus (TAFV), Reston ebolavirus (RESTV), Bundibugyo ebolavirus (BDBV) and Bombali ebolavirus (BOMBV), which have been shown to have different levels of lethality in humans. The genus *Marburgvirus* includes the pathogenic Marburgvirus (MARV). The characteristics of the PE2 of *Filoviridae* have been studied in detail by using ZEBOV as a representative virus. The ZEBOV genome contains PE2 in the NP mRNA 5’ untranslated region (UTR) with uracil (U) at every 6 nts (UN_5_) eight times starting from the 81st nt from 3’ genomic terminus (as shown in Fig. 4) (5,17). Although the significance of having at least three repeated UN_5_ hexamers has been suggested (5), the importance of PE2 in ZEBOV genome replication has not yet been examined in detail. In addition, the knowledge concerning PE2 in other viruses in the family *Filoviridae* remains scarce.

In this study, we comprehensively analyzed PE2 sequences using a public database, and found that almost all paramyxoviruses have C or CG repeats in PE2; the sequence patterns in PE2 were clearly divided in a subfamily-specific manner. The comparable analysis for the position of PE2 and virological properties of the genome replication indicates a unique characteristic specific for the *Rubulavirinae*. We also demonstrated that four repeated UN_5_ hexamers are essential for ZEBOV genome replication. This led to the finding that most viruses in the family *Filoviridae* have at least four repeated UN_5_ hexamers in their genomic and antigenomic promoters.

## Results

### Comparison of all PE2 sequences in the family *Paramyxoviridae*

For the comprehensive analysis of PE2 sequences across paramyxoviruses, complete genome sequences annotated as “*Paramyxoviridae*” were downloaded from the NCBI refseq database, yielding a total of 5,100 sequences (within 10 to 30kb). Within this dataset, 1,647 sequences exhibited the conserved paramyxoviral genome terminus 5’-ACC-GGU-3’, and notably, 99.3% of these sequences conformed to a multiple of six length (Supplementary Table 1). Further classification of the sequences revealed three subfamilies, *Orthoparamyxovirinae*, *Avulavirinae* and *Rubulavirinae*, consisting of 675, 789 and 183 sequences, respectively. To assess sequence conservation, we examined the 73 to 98 nts from the 3’ end, which represent the leader (genomic) PE2, and from the 5’ end, which represent the trailer (antigenomic) PE2, using the WebLogo sequence logo generator (https://weblogo.berkeley.edu/logo.cgi). Almost all viruses in the subfamily *Orthoparamyxovirinae* possess a conserved PE2 sequence pattern characterized by a C at every 6 nts (N_5_C) three times in hexamer numbers (hex#) 14, 15 and 16 within both the leader and trailer PE2 (Fig. 1C). In contrast, in the majority of the *Avulavirinae* and *Rubulavirinae* viruses, CG was found at every 6 nts (CGN_4_) three times in hex# 13, 14 and 15 in both the leader and trailer PE2 (Fig. 1C). While some variation emerged within virus genera (Supplementary Fig. 1), it was clear that PE2 sequence conservation could be definitely categorized into two patterns: N_5_C at hex# 14, 15 and 16 (*Orthoparmyxovirinae*) and CGN_4_ at hex# 13, 14 and 15 (*Avulavirinae* and *Rubulavirinae*).

### Differences in the trailer PE2 location among paramyxovirus subfamilies

The leader PE2 is presumed to reside within the 3’ UTR of the genome, while the trailer PE2 is situated in the 5’ UTR of the genome. However, our previous investigations have revealed that in the human parainfluenza virus type 2 belonging to the subfamily *Rubulavirinae*, the trailer PE2 is localized within the open reading frame (ORF) of viral L protein (18). Consequently, paramyxoviruses may be categorized into two distinct classes: those harboring PE2 outside of the ORF, and those harboring PE2 within the ORF. By examining the length of the 3’ UTR and 5’ UTR of each viral genome, it is possible to determine whether PE2, which is located at 73 to 98 nts from the genomic and antigenomic ends, is situated outside or inside of the ORF. We first comprehensively examined the lengths of the 3’ UTRs in 1647 genome sequences of paramyxoviruses (Supplementary Fig. 1). The shortest 3’ UTR was 98 nts in the Denwin virus in the subfamily *Orthoparamyxovirinae* (GenBank accession number OK623354), indicating that there are no viruses in the database with a leader PE2 inside of the ORF. Subsequently, the lengths of the 5’ UTRs of the genomes were comprehensively examined in the same database. The findings indicated that all viruses belonging to the subfamily *Rubulavirinae*, with the exception of mumps virus, had trailer PE2 sequences embedded within the L ORF (Fig. 2A and B). Furthermore, we observed that the trailer PE2 sequences in only 15 viruses of the subfamily *Orthoparamyxovirinae* were partially within the L ORF (Supplementary Table 1). For all of the other viruses, the trailer PE2 sequences were located outside of the ORF.

**Figure 2.**
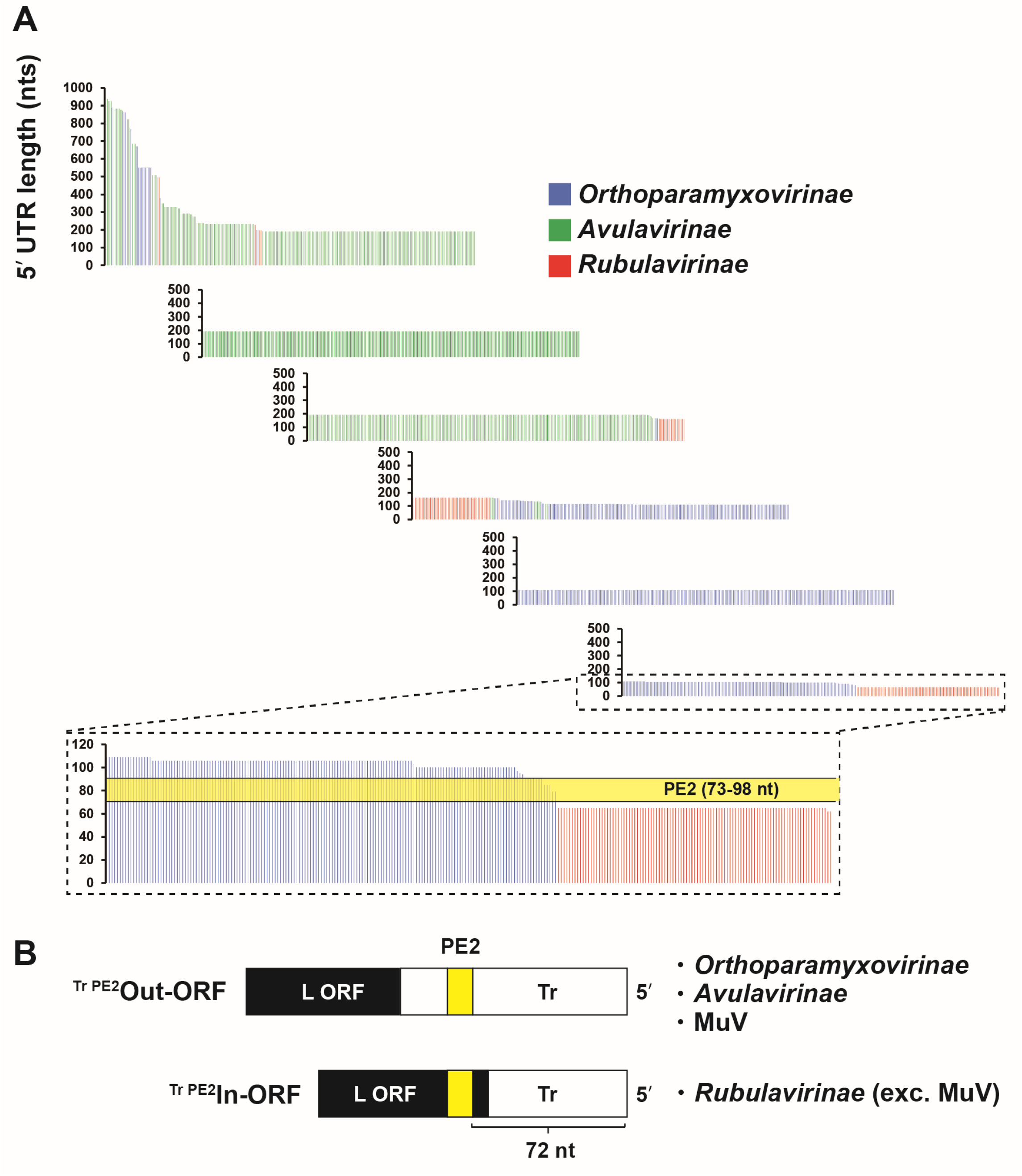
Comprehensive analysis of the genomic 5’ UTR lengths of viruses in the family *Paramyxoviridae.* (A) The nts lengths of the genomic 5’ UTRs of viruses belonging to the subfamilies *Orthoparamyxovirinae*, *Avulavirinae* and *Rubulavirinae*. A bar represents a virus sequence. The trailer PE2 (Tr PE2) region (73 to 98 nts) is shown in yellow. (B) Viruses are divided into Tr PE2 Out- or In-ORF types.

The viruses within the family *Paramyxoviridae* are categorically delineated, as illustrated in Fig. 3A, through a phylogenetic tree predicated upon the alignment of the L proteins (19). The evolutionary trajectory shows an initial divergence of the viruses into an *Orthoparamyxovirinae* lineage and other lineages, which subsequently split into the *Avulavirinae* and *Rubulavirinae* lineages. This primary divergence is strikingly consistent with the divergence of the N_5_C PE2 and CGN_4_ PE2 (Fig. 3A). The *Rubulavirinae* is characterized by the localization of the trailer PE2 within the ORF, and by a different RNA editing mechanism (V-mode) (Fig. 3A). P-mode viruses perform RNA editing to generate the accessory V gene, while V-mode viruses perform RNA editing to generate the essential P gene (Fig. 3B) (20–25).

**Figure 3.**
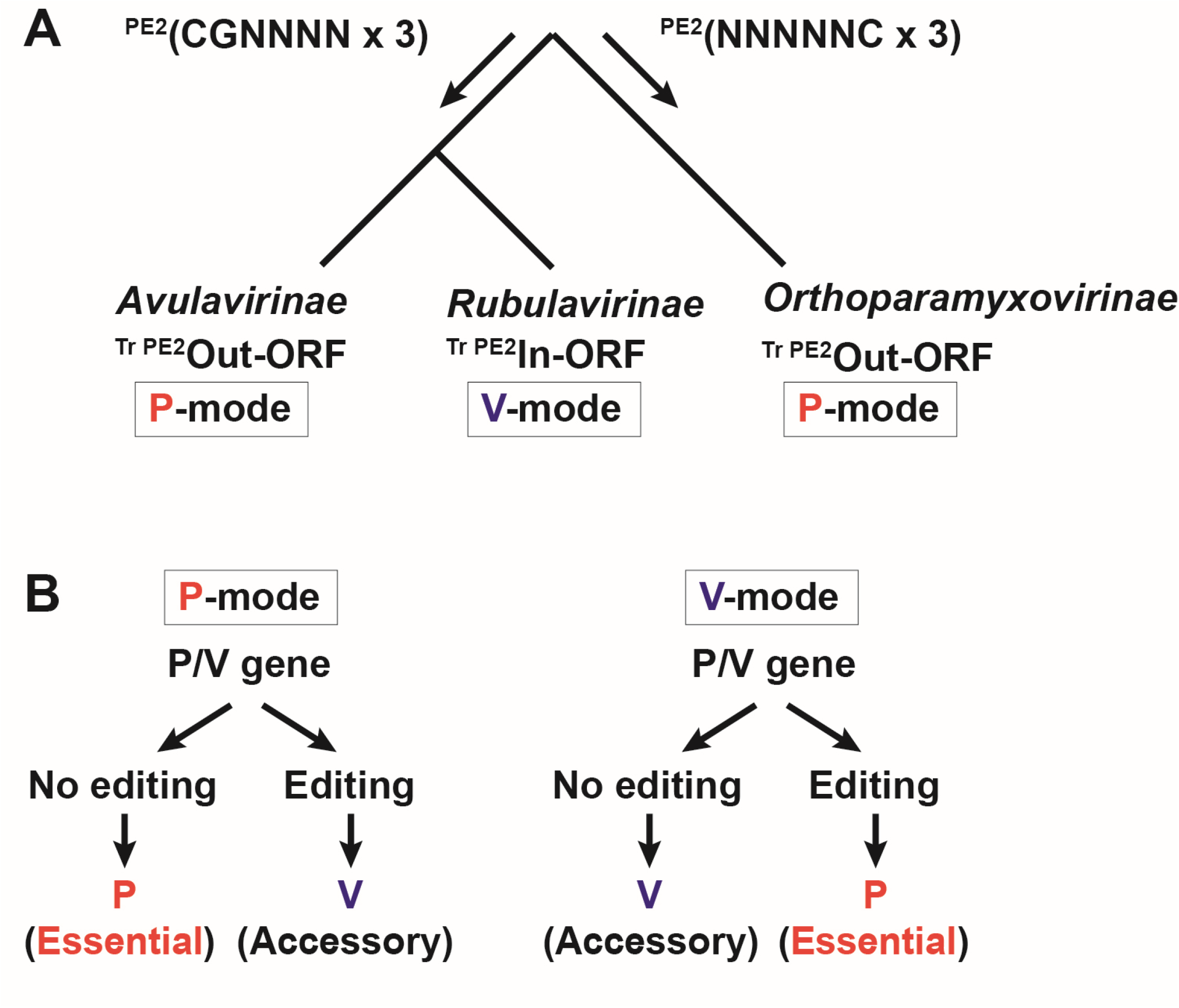
Relationship between PE2 and the virological properties of paramyxoviruses. (A) Relationships among the evolutionary phylogenetic tree, the sequence and location of PE2, and the RNA editing pattern. (B) Differences in the RNA editing modes. The P mode produces an accessory protein via editing, whereas the V mode produces an essential P protein via editing.

### Filoviruses require at least four consecutive UN_5_ repeats for genome replication

Ebolavirus in the family *Filoviridae*, follows the rule of six, which is limited to the promoter portion, and has PE2, similar to paramyxoviruses (5,6). The properties of the PE2 of *Filoviridae* have only been studied in detail in ZEBOV (5,17). ZEBOV has eight consecutive UN_5_, starting at 81 nt from the 3’ end of the genome, and this part is defined as the PE2 (Fig. 4). Using a ZEBOV minigenome assay, it was previously shown that the U at every 6 nts repeated at least three times in this part of the sequence is sufficient for viral genome replication, based on a method of replacing the U with an A (5). However, to date, no study has directly deleted the PE2, and examined the importance of the presence of UN_5_ hexamers in genome replication. Thus, we used the ZEBOV minigenome assay to study the effect of deletion of the entire PE2, *i.e.*, the eight consecutive UN_5_ at 81 to 130 nts of the 3’ genomic end, on genome replication. The results showed that the replication activity was reduced to the same level as the background NP (-), indicating a complete loss of genome replication activity (Fig. 4). Next, the same assay was performed using minigenomes with 6 nts containing a leading U (Uaacuu) added to the deletion region one pair at a time. The results showed that three consecutive occurrences of Uaacuu resulted in the slight recovery of genome replication activity, while four consecutive occurrences of Uaacuu resulted in complete recovery of the activity (Fig. 4).

**Figure 4.**
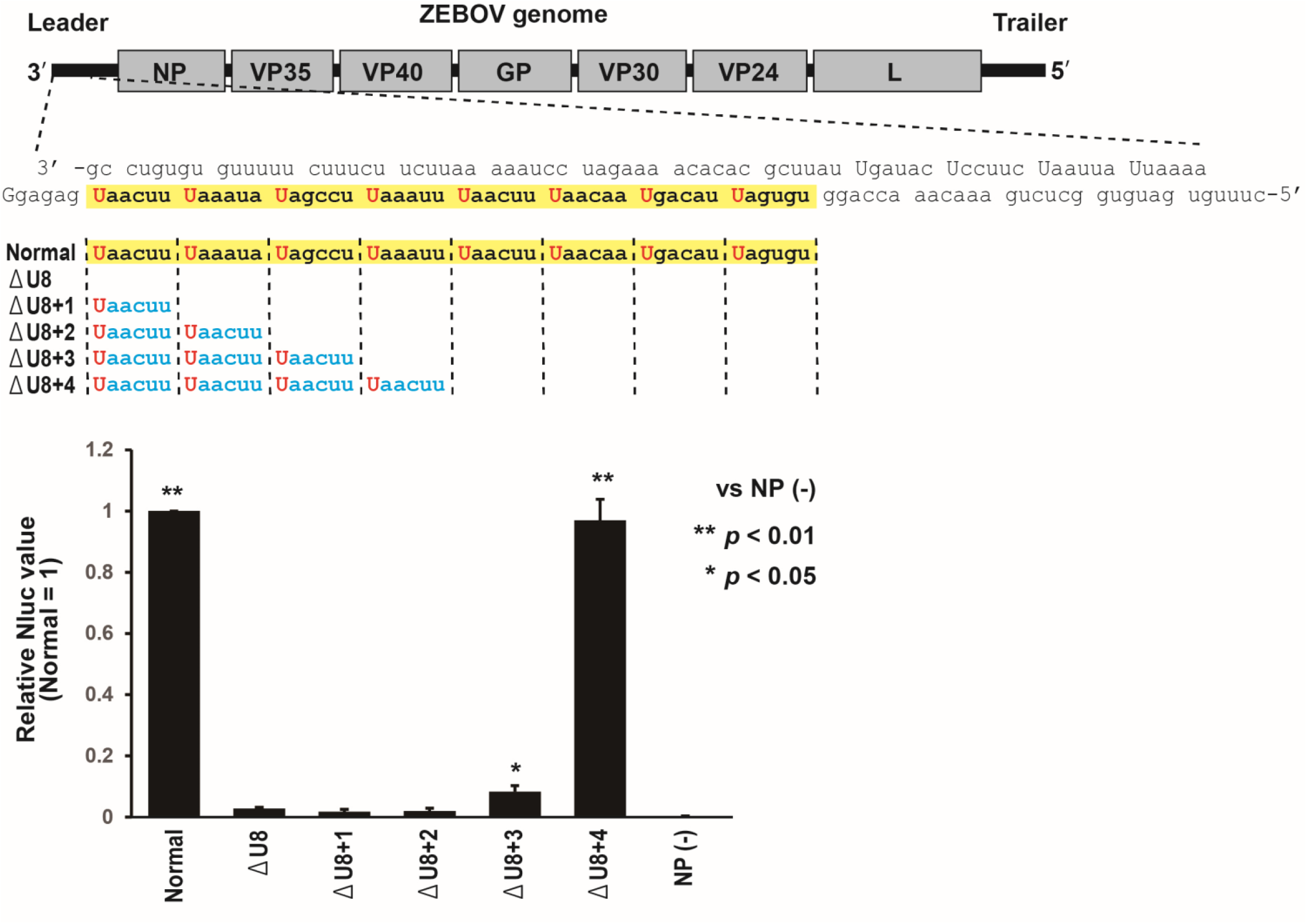
Zaire ebolavirus minigenome study for PE2. ZEBOV PE2 consists of eight sequential UN_5_ repeats. The relative values of Nluc expression are shown with the values of the normal ZEBOV minigenome set to 1. NP (-) indicates the value from the assay using an empty plasmid instead of ZEBOV NP. Bars represent the means and standard deviations (n = 3).

Is the presence of four consecutive UN_5_ a universal trait in all filovirus replication promoters? To answer this question, we searched for regions where UN_5_ appears four or more times at both ends of the genome in all filovirus sequences. Complete genome sequences annotated as “*Filoviridae*” were downloaded from the NCBI refseq database, yielding a total of 4,042 sequences. The viruses in the family *Filoviridae*, unlike the family *Paramyxoviridae*, have an indeterminate terminal nt sequence. Therefore, we extracted genome sequences with the full-genome lengths of 17 to 21 kb. We then searched for sites where U appeared at every 6 nts at least four consecutive times within 200 bases from the end of each genomic sequence. The results revealed that there are sites in the genome and antigenomes of ZEBOV, SUDV, TAFV, RESTV, BDBV, BOMBV, and MARV where consecutive UN_5_ sequences appear four to eight times (Table 1).

**Table 1.**
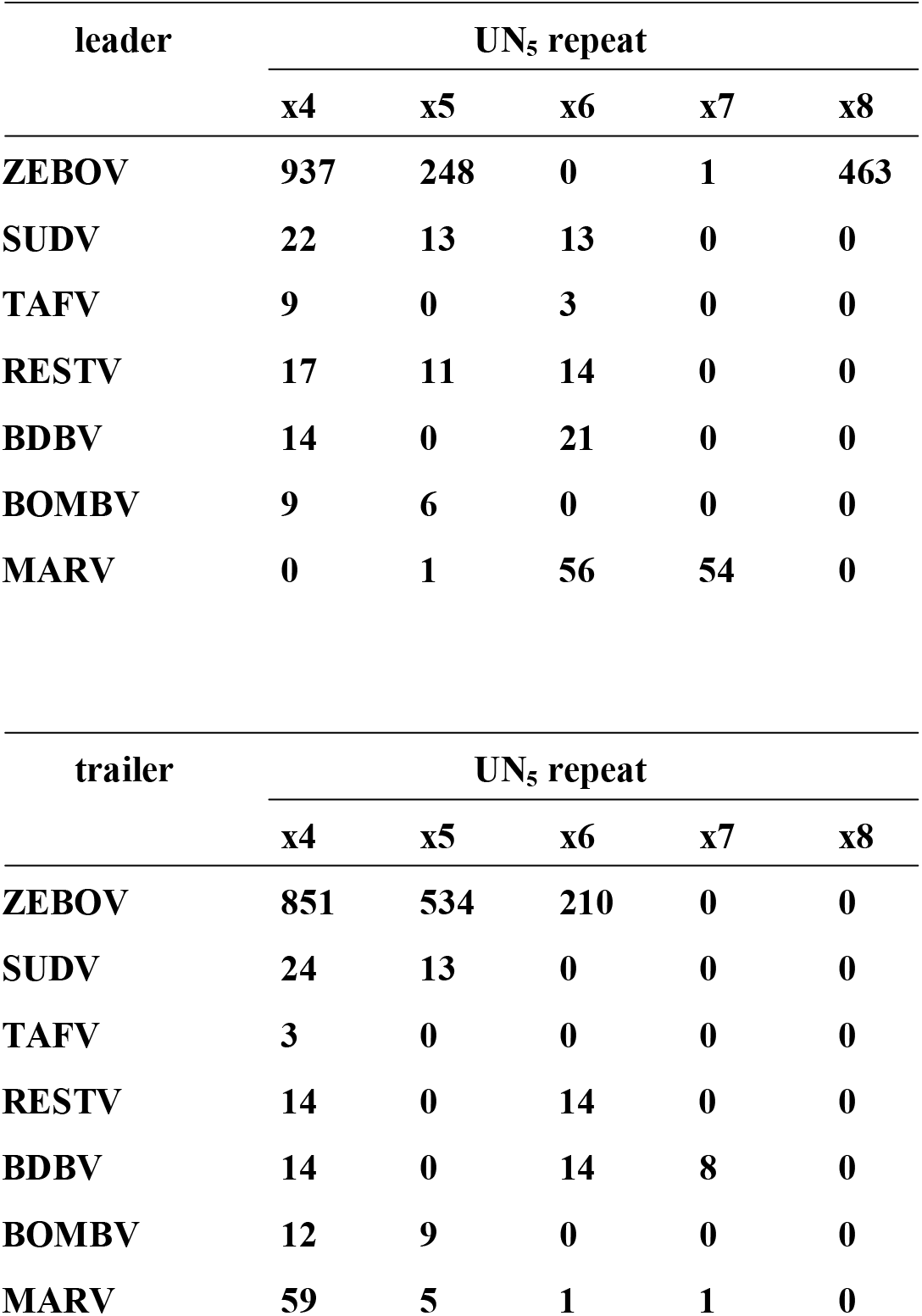
Sequence counts of consecutive UN_5_ hexamer repeats (four to eight times) within 200 bases of the 3’ UTR of the virus genomes (leader) and antigenomes (trailer) in the family *Filoviridae*.

## Discussion

We compared the PE2 sequences, which have only been studied in detail in a limited number of virus species, using a database containing all available paramyxovirus genome sequences. The results showed that the PE2 of paramyxoviruses can clearly be classified into two groups: those with N_5_C at hex# 14, 15 and 16 (*Orthoparmyxovirinae*), and those with CGN_4_ at hex# 13, 14 and 15 (*Avulavirinae* and *Rubulavirinae*) (Fig. 1C). This difference in PE2 pattern is consistent with the evolutionary phylogenetic tree of the viruses (19); the ancestors first diverged into an *Orthoparamyxovirinae* lineage and other lineages and at this point, they also clearly diverged into viruses having PE2 with N_5_C three times or PE2 with CGN_4_ three times. Using the lengths of the 3’ UTR and 5’ UTR as a simple indicator of whether PE2 is located inside or outside of the ORF, we found that all viruses in the subfamily *Rubulavirinae*, except for mumps virus, have the trailer PE2 within the ORF of the L protein (Fig. 2). The *Rubulavirinae* differs from the *Orthoparamyxovirinae* and *Avulavirinae* in that the P mRNA is produced by RNA editing (21,24,25). While the *Rubulavirinae* is strictly reliant on RNA editing for viral propagation, the *Orthoparamyxovirinae* and *Avulavirinae* require it only for accessory protein production (20,22,23). RNA editing efficiency is regulated by an RNA editing signal present on the P gene that functions as a cis-acting element (16), and NPs also bind to this cis-acting element region. Notably, the positioning of the cis-acting element on NPs affects the RNA editing efficiency (26). When NPs bind correctly from the 5’ end of the genome, the cis-acting element is placed correctly on the NPs. The trailer PE2 functions as a promoter during genome replication from the antigenome, and the precise placement of the cis-acting element on NP relies on whether the spacer sequence between the trailer PE1 and trailer PE2 is a multiple of six. Therefore, if there is no mutation in PE2, the insertion/deletion-sensing system of the genome can operate correctly, and the cis-acting element will be placed correctly on the NP. The non-coding region is more susceptible to mutations than the coding region, particularly the ORF for the essential RdRp L protein, which has remained remarkably stable throughout viral evolution. The *Rubulavirinae*, which has a trailer PE2 embedded within the L ORF, appears to have evolved mechanisms to resist insertions, deletions and other mutations in PE2 more effectively than viruses of the other subfamilies. Further research is warranted to determine conclusively whether there is any connection between the RNA editing efficiency and the trailer PE2 sequence.

We also directed our attention to the family *Filoviridae*, as it is another virus group that is subject to the “rule of six” (5,17). Through the deliberate introduction of mutations into the PE2 of the ZEBOV minigenome, we ascertained that the presence of Us at intervals of 6 nts at least four times is imperative for the full minigenome replication of ZEBOV (Fig. 4). Weik et al., have shown that three consecutive UN_5_ hexamers located in the proper phase in the PE2 were sufficient for ZEBOV replication activity (5). Although they suggested that replication occurs more efficiently when more hexamers are present, our minigenome experiment revealed that replication activity peaks when there are four consecutive hexamers. This requirement for UN_5_ hexamers to occur four or more times was found within the 200-bases of both the genome and antigenome termini in the *Filoviridae* (Table 1). The leader PE2 exhibits a greater number of UN_5_ hexamer repeats in comparison to the trailer PE2; in particular, there were eight and seven repeats in the leader PE2 of the ZEBOV and MARV genomes, respectively (Table 1). Longer repeats of UN_5_ may help ensure that four or more contiguous hexamers remain even if a mutation occurs in one of the UN_5_ sequences, and may contribute to maintaining the ability to replicate the genome. It is possible that a correlation exists between the extent of UN_5_ continuity and viral pathogenicity. In fact, only ZEBOV and MARV have ever recorded an approximately 90% case fatality rate among viruses in the family *Filoviridae* (27,28).

Viruses employing an RNA editing mechanism during mRNA transcription concurrently have bipartite promoters. Paramyxovirus RdRp recognizes the contiguous C sequence within the RNA editing site on the P gene as a cis-acting element, and inserts G(s) into the mRNA during the transcription process. Filovirus RdRp recognizes the contiguous U sequence within the RNA editing site on the glycoprotein gene, and inserts A(s) into the mRNA. The PE2 of paramyxoviruses and filoviruses comprises C and U repeats at every 6 bases, respectively. Thus, there appears to be a mechanism whereby paramyxovirus RdRp recognizes C-based elements and filovirus RdRp recognizes U-based elements.

## Materials and Methods

### Cells

Baby hamster kidney (BHK) cells constitutively expressing T7 RNA polymerase (BHK/T7-9 cells) (29) were cultured in Dulbecco Modified Eagle’s Medium with 5% fetal calf serum and penicillin/streptomycin. Cells were cultured at 37°C in 5% CO_2_.

### Plasmid construction

The nanoluciferase (Nluc)-expressing ZEBOV (GenBank accession number AF086833) minigenome plasmid (ZEBOV-Nluc) was constructed using the pUC57 plasmid backbone. The Nluc gene is flanked by the 3’ UTR (469 nts) containing the leader sequence and the 5’ UTR (740 nts) containing the trailer sequence of the ZEBOV genome. The minigenome is set under the control of the T7 RNA polymerase promoter, and the transcript expressed as a negative-sense RNA is cleaved at both ends by a hammerhead ribozyme and a hepatitis delta virus ribozyme (30). The ZEBOV NP, VP35, VP30 and L genes cloned into a pCAGGS vector were as described previously (31). pCAGGS firefly luciferase (Fluc) was also constructed. The deletions of the PE2 in ZEBOV-Nluc were performed by a standard cloning method.

### ZEBOV Nluc minigenome assay

The ZEBOV Nluc minigenome assay was performed in BHK/T7-9 cells cultured in 12-well plates. Plasmids; ZEBOV-Nluc (0.5 μg), pCAGGS-L (0.4 μg), -VP35 (0.4 μg), -VP30 (0.4 μg) and -NP (0.4 μg) or empty vector, and Fluc (0.05 μg) were transfected using XtremeGENE HP (Merck, Darmstadt, Germany). At 48 h post-transfection, the Nluc and Fluc activities were measured using the Nano-Glo Dual-Luciferase Reporter Assay System (Promega, Madison, WI, USA) according to the manufacturer’s instructions. All obtained results of Nluc were normalized by the expression levels of Fluc.

### Analysis of the PE2 sequences and UTR lengths of the viruses in the family Paramyxoviridae

Sequences annotated as *Paramyxoviridae* were downloaded from the NCBI refseq database on August 3rd, 2022. There were 64,057 sequences annotated as *Paramyxoviridae* in the database. Sequences shorter than 10,000 nt and longer than 30,000 nt (n = 58,957) were excluded from the analysis. Within the obtained dataset, 1,647 sequences exhibited the conserved paramyxoviral genome terminus 5’-ACC-GGU-3’, and 99.3% of these sequences conformed to a multiple of six length. For each sequence, we detected the genome length, 3’ UTR length, 5’ UTR length, sequences of the trailer PE2 (73 to 96 nts from 5’ terminus of the genome), and sequences of the leader PE2 (73 to 96 nts from the 3’ terminus of the genome) (Supplementary Table 1). The codes used for this analysis are available on GitHub (https://github.com/shohei-kojima/Filo_Paramyxo_2023).

### Search for UN_5_ repeats in the genome sequences of the viruses in the family *Filoviridae*

Sequences annotated as *Filoviridae* were downloaded from the NCBI refseq database on July 26th, 2023. There were 4,042 sequences annotated as *Filoviridae* in the database. Sequences shorter than 17,000 nt and longer than 21,000 nt (n = 601) were excluded from the analysis. The presence of UN_5_ repeats was detected by a custom Python script which detects four or more consecutive units of UN_5_ within the 200 nt from the ends of the input sequences (Supplementary Table 2). The codes used for this analysis are available on GitHub (https://github.com/shohei-kojima/Filo_Paramyxo_2023).

### Statistical analysis

Statistical analyses were performed with Prism software (version 9.1.2; GraphPad, San Diego, CA, USA). Statistical significance was assigned when p values were <0.05. Inferential statistical analysis was performed by one-way analysis of variance followed by Tukey’s test.

### Availability of data

All databases used in this study are available from DDBJ/ENA/GenBank (https://www.ddbj.nig.ac.jp/about/insdc-e.html). The accession numbers of the viral sequences used in this study are listed in Supplementary Tables 1 and 2.

## Acknowledgements

We thank Dr. Naoto Ito (Gifu University) for providing the BHK/T7-9 cells, and Dr. Thomas Hoenen (Friedrich-Loeffler-Institut) for providing the plasmids encoding ZEBOV genes. We thank Dr. Kaoru Takeuchi (University of Tsukuba), Dr. Akatsuki Saito (University of Miyazaki), Dr. Makoto Ozawa (Kagoshima University), Dr. Tetsuya Tanaka (Kagoshima University) and Dr. Michinori Kohara (Tokyo Metropolitan Institute of Medical Science) for providing the equipment for biological experiments. We also thank Dr. Kyoko Tsukiyama-Kohara (Kagoshima University) and members of her laboratory for their support with the biological experiments.

---

**Supplementary Figure 1.**
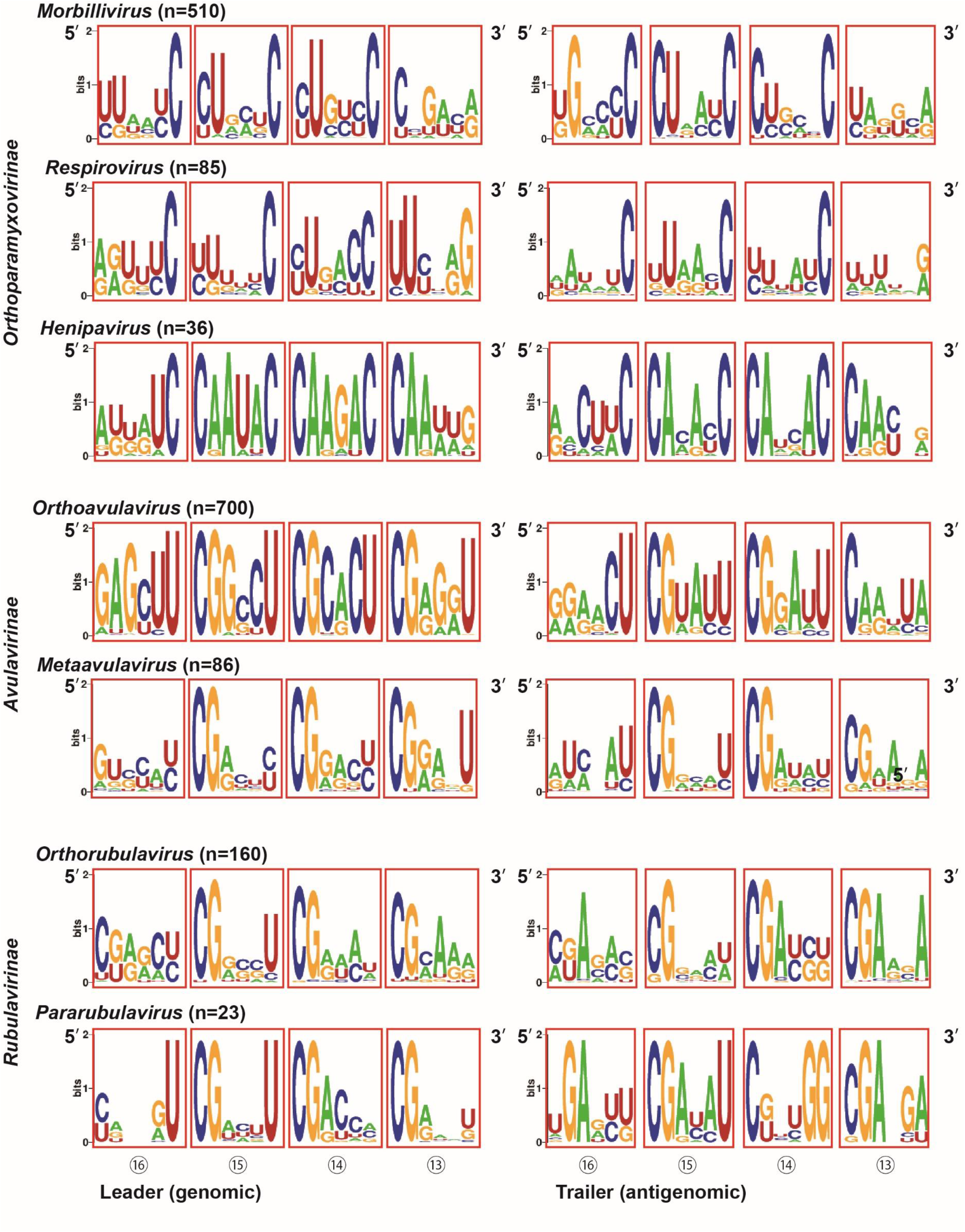
Extensive comparative analysis of the PE2 sequences of each virus genus in the family *Paramyxoviridae*. Conserved nts within the leader and trailer promoters of viruses belonging to the genus *Morbillivirus, Respirovirus* and *Henipavirus* (subfamily *Orthoparamyxovirinae*), *Orthoavulavirus* and *Metaavulavirus* (subfamily *Avulavirinae*), and *Orthorubulavirus* and *Pararubulavirus* (subfamily *Rubulavirinae*). The numbers in the circles on the diagram indicate the number of the hexamer from the 3’ terminus. Each red square indicates a NP hexamer.

**Supplementary Figure 2.**
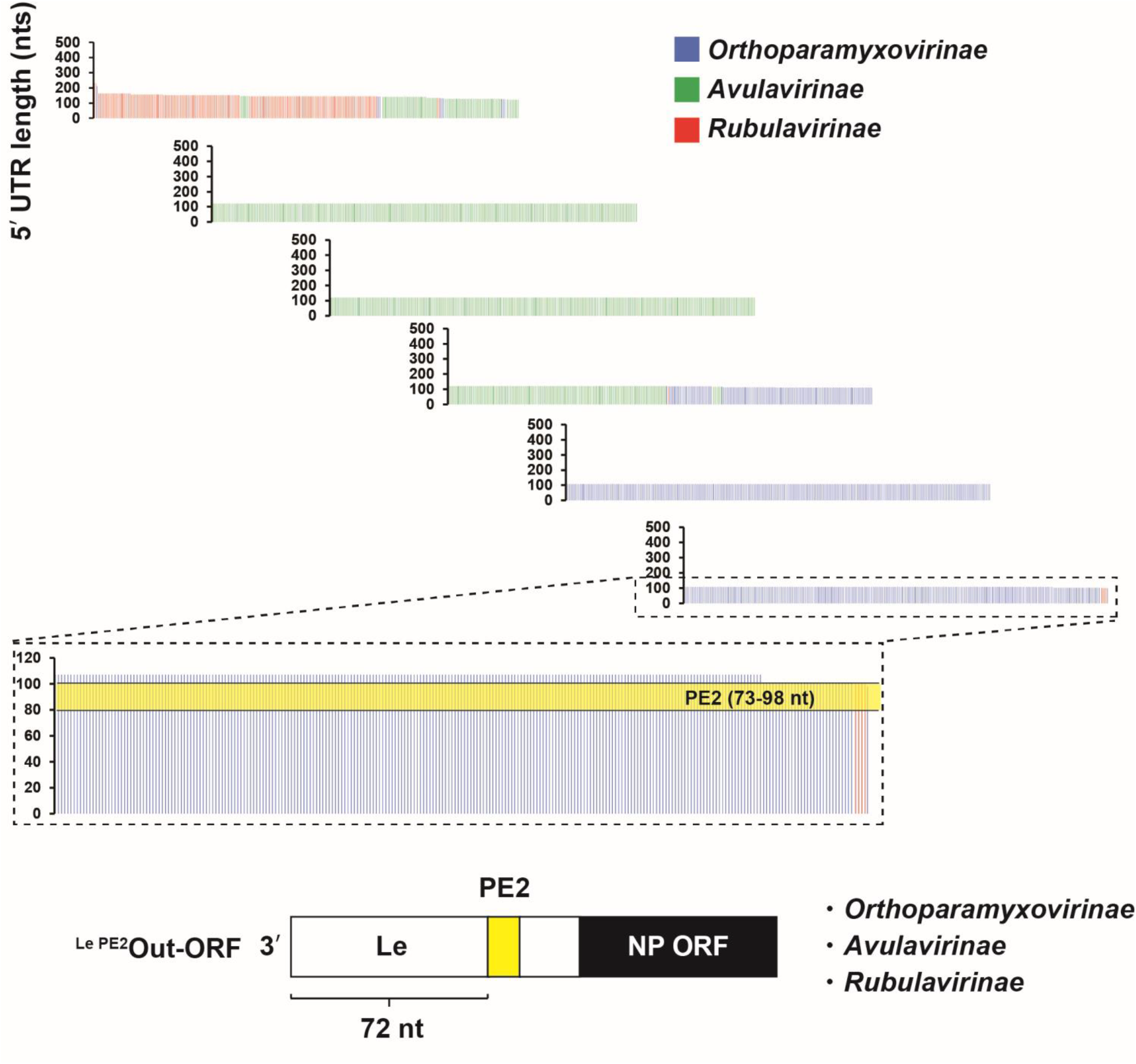
Comprehensive analysis of the genomic 3’ UTR lengths of viruses in the family *Paramyxoviridae*. The nts lengths of the genomic 3’ UTR of viruses belonging to the subfamilies *Orthoparamyxovirinae*, *Avulavirinae* and *Rubulavirinae*. A bar represents a virus sequence. The leader PE2 (Le PE2) region (73 to 98 nts) are shown in yellow.

